# Particulate matter concentration in Davao City airshed and its trend over the years

**DOI:** 10.1101/2023.07.27.550832

**Authors:** Marife B. Anunciado

## Abstract

Davao City airshed was selected for air quality mapping using particulate matter (PM) concentrations. PM data were taken from the regulatory office, Environmental Management Bureau XI, from 2016 to 2021 to understand annual variation and determine trends that may be attributed to seasonal changes in the region. PM concentrations were spatially interpolated using Inverse distance weighting (IDW) feature, an interpolation technique of ArcGIS. PM concentration and distribution over the years showed no similar patterns, both for PM_10_ and PM_2.5_. No annual similarities of PM concentration were observed, and distribution varies yearly. No seasonal trends were shown on the interpolated maps for PM. However, there was an overall PM concentration decrease and distribution covered fewer affected areas over time. PM concentration in 2016 were generally at a level within the defined limit of NAAQGV except for some AQMS locations and years but sparingly exceeding the NAAQGV limit over time. Results show that PM emissions were lower suggesting a possible success on the regulation policies in the Davao City airshed through reduction or better management of air pollutant emissions.

## 1. Introduction

Air pollution is an environmental problem that is continually increasing and impacts more countries over time. While some countries, mostly developed nations have successfully managed air pollutant emissions into the air, more nations must extensively prioritize management of air pollution. A global scale study identified a poor performance on reducing exposure to outdoor pollution by countries and continuing efforts should be strengthened to lessen the risk of public to air pollution ^1^. Air pollution source can be anthropogenic or may be a result of natural process (e.g., soil dust, sea salt, carbon, biogenic) containing organic matter, sulfates, and carbonaceous emission ^2^. Various air pollutants contribute to the overall quality of ambient air such as oxides (sulfur, nitrogen, carbon) and particulate matter (PM). Relative to SO_2_ and NO_2_, PM_10_ concentration in Hyderabad, India was extremely higher than the WHO prescribed values to more than 3 order consistently in years (2016-2018) ^3^. PM pollution can become a serious environmental problem especially when regions suffer from a dry climatic condition ^4^.

PM refers to any chemical, physical, and biological substances that exist as particles in many shapes and sizes ^5^. Particle size affects the transportation and deposition of PM in the respiratory system, thus, is directly important and relevant in determining health effects upon exposure to PM. PM emission ranks with the highest increase in risk exposure from 1990 to 2019 globally. Deaths attributed to outdoor PM pollution accounts to 7.8% of global deaths ^1^. Philippines, according to a global systematic analysis, suffer from air pollution where ambient PM, ambient ozone and household indoor pollution all contributed in the same manner ^1^. Air pollution level in the country led to an 8-10% disability-adjusted life-years. Public utility jeepneys, a common public transportation vehicle in Philippines, emitted significantly higher (60% more) black carbon than lighter-duty vehicles (LDVs) in Manila, Philippines ^6^. Furthermore, deaths due to outdoor air pollution accounts to 4% in 1990 and increased to 5% in 2019 ^7^ in the country. For a given 100,000 individuals, about 36 of those deaths in 1990 and 44 in 2019 were influenced by exposure to air pollutants and could be more potentially devastating as more people are exposed to outdoor pollution due to daily outdoor engagement and responsibilities ^8^. An exposure to PM_2.5_ of a city in Philippines were majorly contributed by vehicle emissions, household emissions from cooking and burning of agricultural waste ^9^.

Air quality monitoring stations (AQMS) provides information on air quality levels but could only be specific and restricted to nearby locations of AQMS at a particular time. This does not guarantee AQ concentration uniformity on a wider scale. Mapping of air pollution is one approach of monitoring air pollution and prediction of monitored pollution level provided reliable estimates ^10^. Mapping of air quality (AQ) through GIS was utilized to evaluate the cost of health care of exposed individuals to various air pollutants estimating economic costs due to health damage ^11^, characterizing elemental constituents of dust ^4^ and understanding land cover and human activities impacts to air pollutant emission ^9^. The IDW technique was utilized in various air pollution studies ^3,11–13^. Mapping air pollution concentration aid difficulty in air pollution monitoring and may assist on the identification of hotspots for air quality management needs across the city.

In Davao City, intensifying ambient air quality management efforts by establishing more sampling stations to protect public health is a priority ^14^. This study aims to characterize the variability of air quality in the Davao City airshed using an interpolation technique in ArcGIS.

The criteria pollutants cover PM concentrations, PM_10_ and PM _2.5_. Specific objectives were to 1) evaluate PM_10_ characteristics, seasonally and annually and 2) evaluate PM_2.5_ characteristics on annual basis. Output maps will be helpful in assisting future decision making and aid in formulation of air regulatory policies in the Davao City airshed.

## 2. Materials and Methods

### 2.1. Study area

Davao City, situated in the southeast region of Philippines, covers a 244,000-hectare land comprised of residential communities, commercial establishment, and agricultural areas. Registered vehicles account at 170,000 and Davao City can expect 586,000 vehicles accessing the city routes on daily regular basis. As the central trade in the south, 544 industrial firms operate in the region with majority of food manufacturing, trade and commercial facilities ^15^. Air quality of Davao City is regularly monitored by the Environmental Management Bureau Region XI of the Department of Environment and Natural Resources. EMB XI have established six air quality monitoring stations (AQMS) in the Davao City airshed, mostly along coastal regions where the hub of urban development is situated. The AQMS were constructed strategically located where mobile and area sources of air pollutants is expected to be high and in the same part of the region where population density is also high (Figure 1). The AQMS are located in 1) Bugac, Ilang; 2) Barangay 12-B; 3) Davao Memorial Park; 4) Toril Plaza; 5. Calinan National High School PMS; and 6) Davao International Airport. Among the programs promoted in the city to promote better air quality includes construction of city islands as a suspension management for dust, rehabilitation of urban greenery and forested parks as part of the nationwide establishment of carbon sink areas in the country and the anti-smoke belching ordinance.

**Figure 1.**
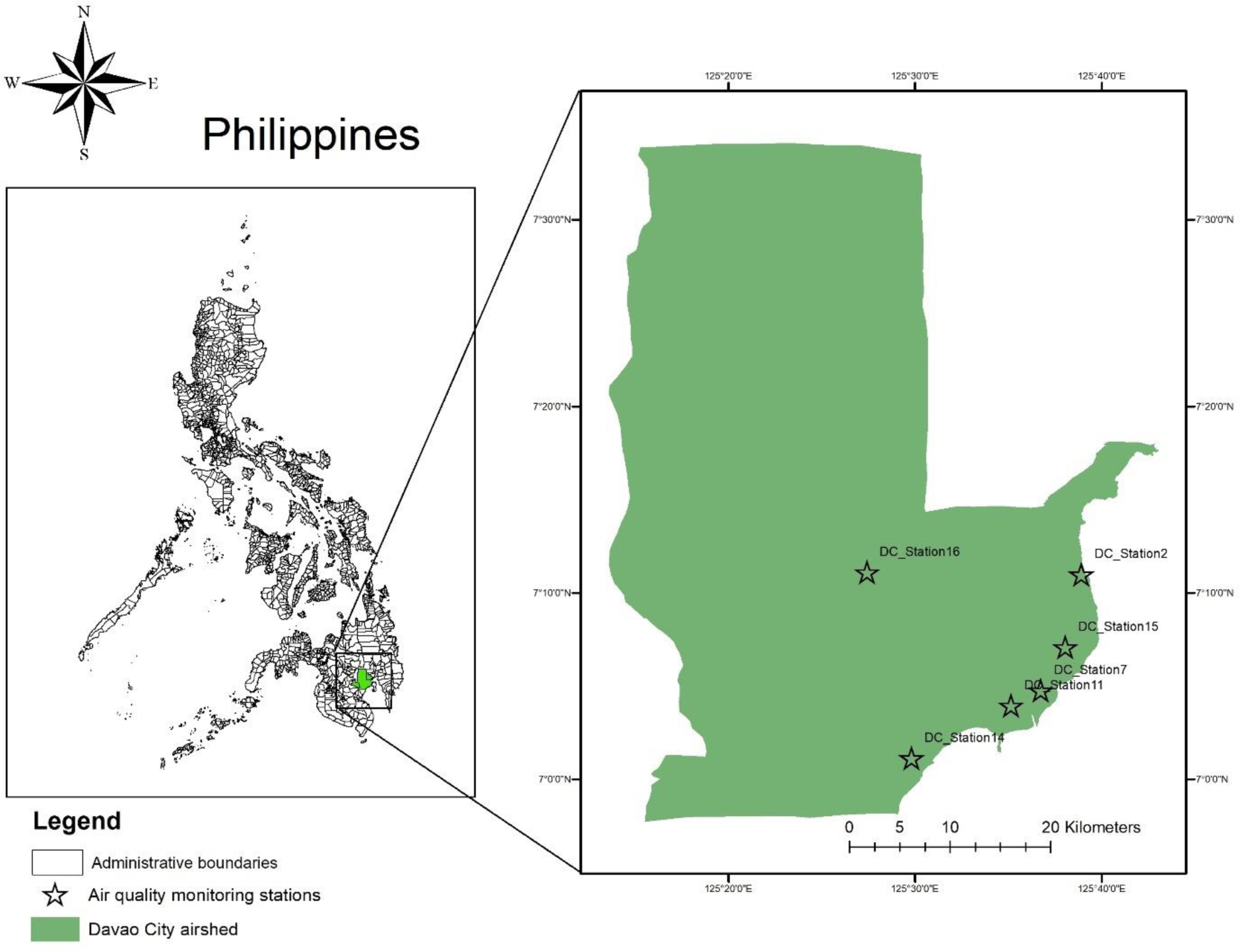
The Davao City airshed and the six air quality monitoring stations (AQMS) providing air pollutant values along with other air quality parameters.

The climate of Philippines can be divided into two major seasons: (1) the rainy season, from June to November; and (2) the dry season, from December to May. The dry season may be subdivided further into (a) the cool dry season, from December ^16^. In 2005, total rainfall was recorded at 1,499 mm during the dry months from February to April and wet months from May to January ^15^.

### 2.2. Air pollutant: Particulate matter

Particulate matter (PM_10_ and PM2.5) is among the six criteria pollutants regularly monitored and prioritized by the Philippine Clean Air Act of 1999 (RA 8749) to reduce air pollution and ensure air quality health parameters in the country ^17^. Particulate matter, specifically PM2.5, reaching the lower respiratory tract can critically impact human health. Characterizing the particulate matter concentration and distribution in the Davao City airshed will contribute to understanding the risks exposure to PM and will aid us in finding a solution to monitor and mitigate what is locally the pressing situation in the region.

### 2.3. Spatial analysis and Data Source

Air quality data (e.g., PM concentration), geographic coordinates of AQMS and shapefile of Davao City (e.g. geographic attributes) were essential parameters needed to develop air quality concentration maps. Geographic attributes of Davao City was taken from Earthworks of the Stanford Libraries ^18^.

#### Data Source

The air quality data, specifically particulate matter concentrations, were taken from 6 AQMS in the Davao City airshed governed by the DENR-EMB XI located in Guzman St., Brgy 27-C, Davao City. EMB XI supervise the operation of these stations as well as delivers the information and cautionary measures to the public in collaboration with the Davao City Air Governing Board in support to EMB XI’s mandate to disseminate information and awareness on health-related issues due to air pollution.

#### IDW

Particulate matter concentrations are utilized into the geospatial database and accessed for complex manipulation; for this study, PM concentrations were explored using the ArcGIS Geostatistical Analyst feature to create surfaces reflecting PM concentration and distribution in the Davao City airshed. All PM concentrations in microgram per normal cubic meter (ug/Ncm) were collected and supplemented by EMB XI. Mean concentrations from January to December were determined. Annual mean concentrations from 2016-2021 were used as input parameters in ArcGIS to generate maps that reflects spatial distribution of PM concentrations.

## 3. Results and discussions

Particulate matter (PM) _10_ or inhalable particles are particle sizes equal to _10_ micrometers in diameter or smaller but not smaller than 2.5 micrometer. PM_10_ concentrations (ug/Ncm) used for spatial interpolation using ArcGIS characterized the air quality condition in the Davao City airshed from 2017 to 2021 (Figure 2). PM_10_ concentration in the region underwent a clear decrease of PM_10_ in six years. Few critical areas (56-60 ug/Ncm), areas where PM_10_ concentration was close to reaching the defined limit for PM_10_, were identified near two (2) AQMS. Close to these two stations exists a school (Calinan National High School) and a commercial establishment (Toril Plaza) may influence PM_10_ emission sources. In India, PM_10_ had a major contribution relative to NO_2_ and SO_2_ to the overall AQI level as concentration continuously increase in areas with high industrial activities and where educational institutions were established ^19^. PM_10_ concentration of a central business district in the country significantly contribute to the local surface dust measured ^20^. Further identification of PM_10_ sources were known to be originating from soil sources, road-dust re-suspension, and vehicle emissions. It is then important to know how local activities can greatly and directly impact PM levels of an area. While it is shown that PM_10_ in 2017 was relatively higher compared to other years, PM_10_ concentration was within the defined limits established by the National Ambient Air Quality Guideline Value (NNAAQGV) of the Philippine Clean Air Act of 1999 at PM_10_ – 80 ug/Ncm annual mean. PM_10_ distribution varies a little bit in 2018 than 2017 where only fewer areas were identified critical to high exposure to PM_10_ but close to the same station where the Toril Plaza is located. PM_10_ in 2019 looks similar and do not vary with the years onward (2020-2022) which clearly shows a decreasing trend of PM_10_ concentration in the Davao City airshed. Overall, PM_10_ distribution varies yearly, and no similar pattern was observed over the years.

**Figure 2.**
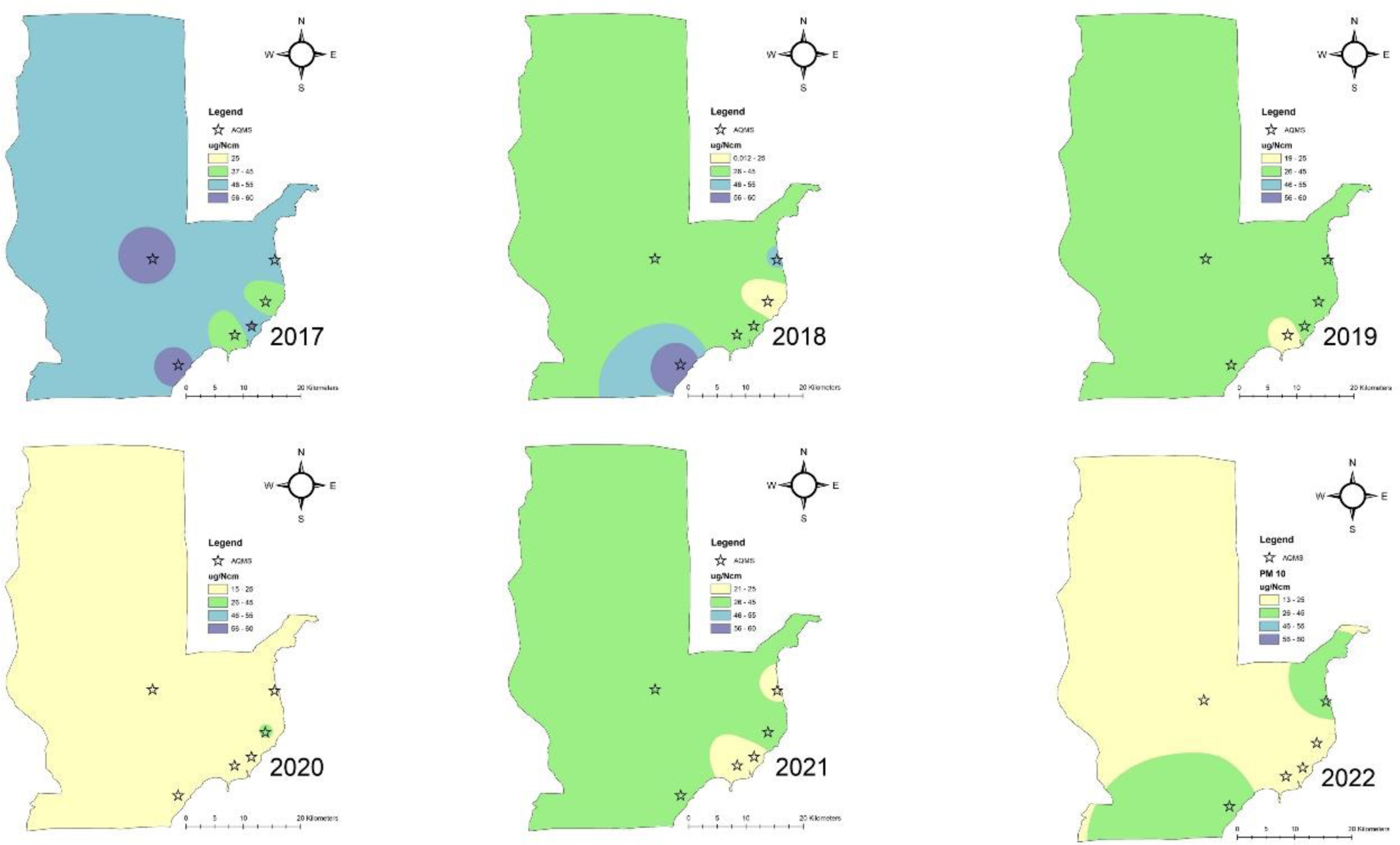
Particulate matter (PM_10_) concentration in the Davao City airshed using the mean concentration of each respective year from 2017 to 2022 (ug/Ncm).

Studies from other countries showed an increasing trend of PM_10_ concentration with values ranging from a minimum of 61 ug/m^3^ to 182 ug/m^3^ in 2016 to 2018 ^3^, relatively higher than the concentration in the Davao City airshed. Significant decreasing trend of PM_10_ was observed from 2005 to 2017 in Spain, which is within the defined limit of European Union but not World Health Organization (WHO) limit ^21^, values also higher than the study. The economic crisis impacts economic activity leading to unemployment and could possibly the reason of reduction of anthropogenic emissions of PM_10_. In China, areas in the coastlines were shown to have better air conditions (lower AQI) than the rest of the regions which can be attributed to better air circulation on areas near the coast than the areas farther from the coast ^22^. This may probably explain the low concentration levels in the AQMS in the city and could have potentially impact PM_10_ concentration levels in the region. A descriptive and indirect way of PM characterization is using the Air Quality Index (AQI). AQI levels ranges from a good to fair, unhealthy to emergency categories which are linked to human health and individual’s susceptibility to respiratory issues upon exposure to an air pollutant ^14^.

PM_2.5_ concentrations (ug/Ncm) was also used for spatial interpolation using ArcGIS to understand the status of air quality in the Davao City airshed. Characterizing PM_2.5_ other than PM_10_ information further strengthen air quality needs and management in the region. However, unlike PM_10_, PM_2.5_ measurement capability was only performed in two (2) AQMS’, namely DC Stations 15 (Davao International Airport) and 16 (Calinan National High School). These two stations were employed with automatic sampler with a continuous sampling frequency. PM_2.5_ has a prescribed annual mean limit of 25 ug/Ncm by NAAQGV. As shown in Figure 3, PM_2.5_ concentration in the Davao City airshed is on a high level but only on areas closer to the two AQMS, particularly to DC Station 16 near the national high school. While other areas are clearly within the defined limits for PM_2.5_ of NAAQGV, it is important to know that spatial interpolation may be underestimated due to lack of PM_2.5_ data from other AQMS in the Davao City airshed. Regardless of this missing information, PM_2.5_ distribution varies year to year. Figure 3 shows that PM_2.5_ concentration was improving over three (3) years from 2017 to 2019. The areas around DC Station 16 initially covered more areas in 2017, had grown smaller by 2019 and eventually disappeared by 2020 until 2021 until no further critical areas were identified. There were critical areas identified around DC Station 15 on two years (2017-2018) which slightly expanded in one year from 2017 to 2018 (Figure 4). In 2019 until recent year of study (2021), PM_2.5_ concentration decreased and remained at a level within the defined limit of NAAQGV. Overall, PM_2.5_ distribution varies yearly, and no similar pattern was observed over the years. Seasonal variability (2016-2021) and its effect to PM_2.5_ distribution was not performed due to limited PM_2.5_ data.

**Figure 3.**
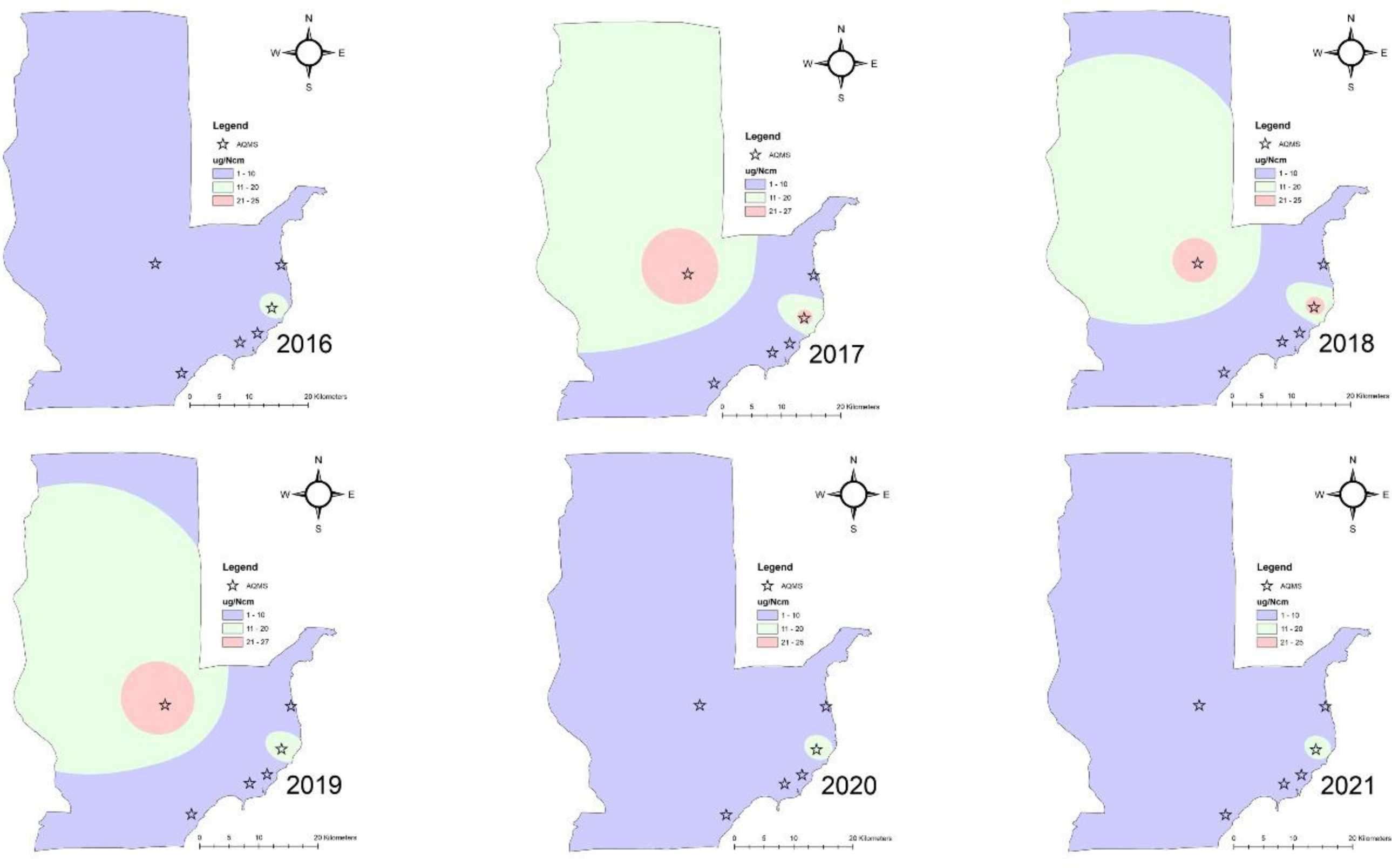
Particulate matter (PM_2.5_) concentration in the Davao City airshed using the mean concentration of each respective year from 2016 to 2021 (ug/Ncm).

**Figure 4.**
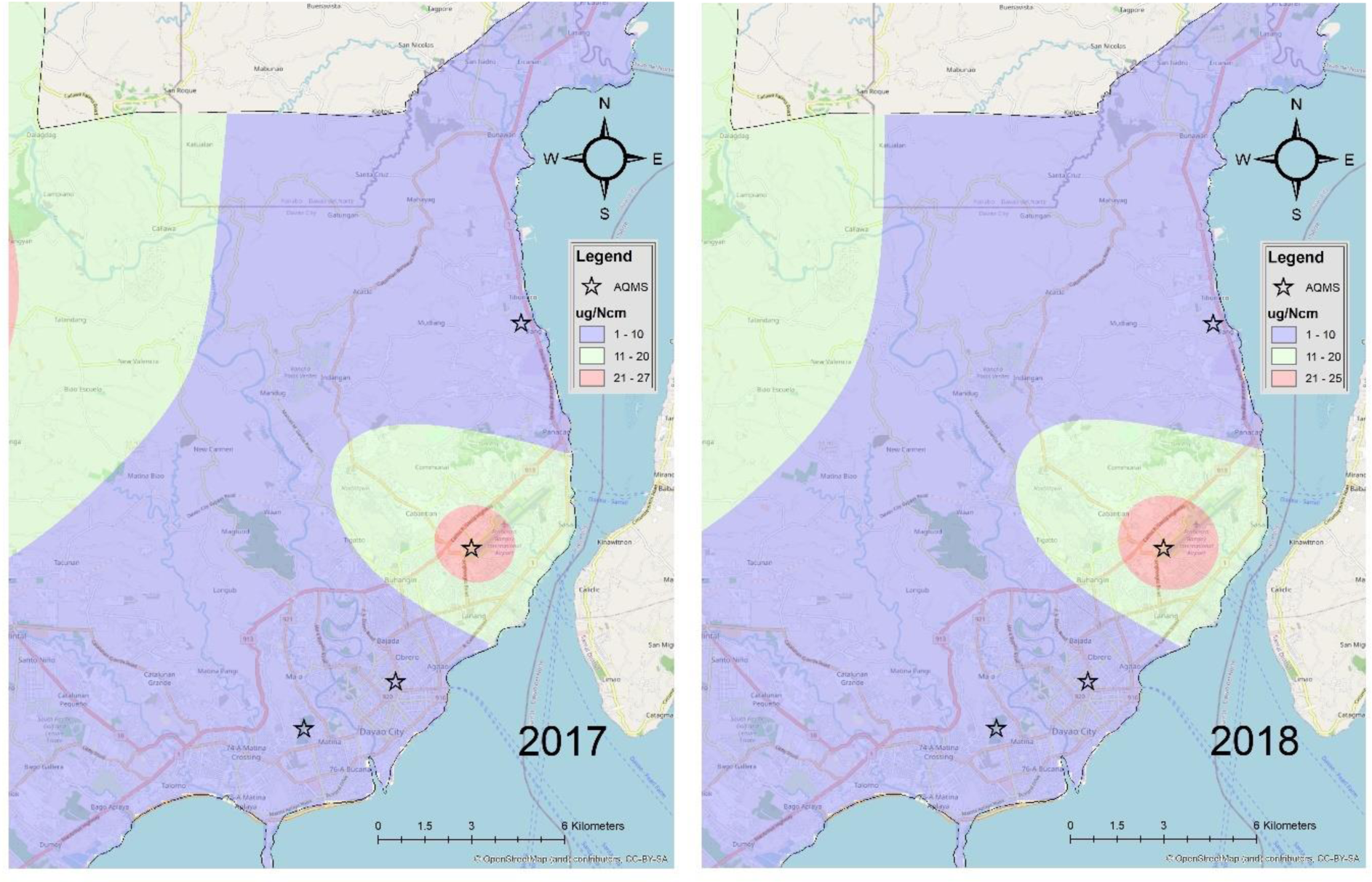
Particulate matter (PM_2.5_) concentration (ug/Ncm) in the Davao City airshed in 2017 relative to 2018.

PM_10_ distribution and condition in our study, however, do not show similar results to an air quality study conducted (2016-2020) in the National Capital Region (NCR) of Philippines using the similar interpolation technique. The AQI in NCR relative to PM_10_ concentration in Davao City airshed showed an overall good level of air quality in Metro Manila and observed to be more prevalent during wet season than dry season. Dry season (December-May) tends to have a moderate AQI level whereas wet season (June-November) have moderate to good AQI ^12^. On an opposite situation, a city in China had a good air quality (lower AQI) during summer than winter season ^22^. In a study using mapping of AQI, season (winter vs. summer) did not vary and also resulted to a similar trend despite of rainfall variabilities ^19^. Factors affecting PM_10_ concentration and distribution can be influenced by factors primarily with air pollution sources and intensity of emission as well as meteorological condition (e.g. rainfall, wind speed, wind direction). The climate of the Philippines was categorized into four characteristics according to occurrence of dry and wet season. In a region where PM concentration (e.g. dust) was massive in a year (12,057 tons/2015), PM distribution had higher PM levels in the spring (200 g/m^2^) and lower levels in the fall (200 g/m^2^) ^4^. Among the factors affecting the high PM were higher building density, less rain and increased wind erosion upon interpreting generated output maps of air quality in the area.

Figures 5, 6, 7, 8 and 9 shows the monthly average PM_10_ concentration in the Davao City airshed from 2017 to 2021. PM_10_ annual limit was utilized as a guideline because there was no prescribed monthly long-term monitoring limit for PM_10_. Few months in 2020 do not show PM_10_ concentration due to unavailable data. Results show no PM_10_ concentration trend over the years (2017-2021) and no seasonal influence observed. The Davao City airshed falls into the 4^th^ category of climatic condition wherein rainfall has more or less distribution throughout the year (DOST-PAGASA) compared to wet season of NCR from June to September. This could possibly explain the opposite result and absence of distinguishable difference on PM_10_ concentration between dry and wet season in the Davao City airshed.

**Figure 5.**
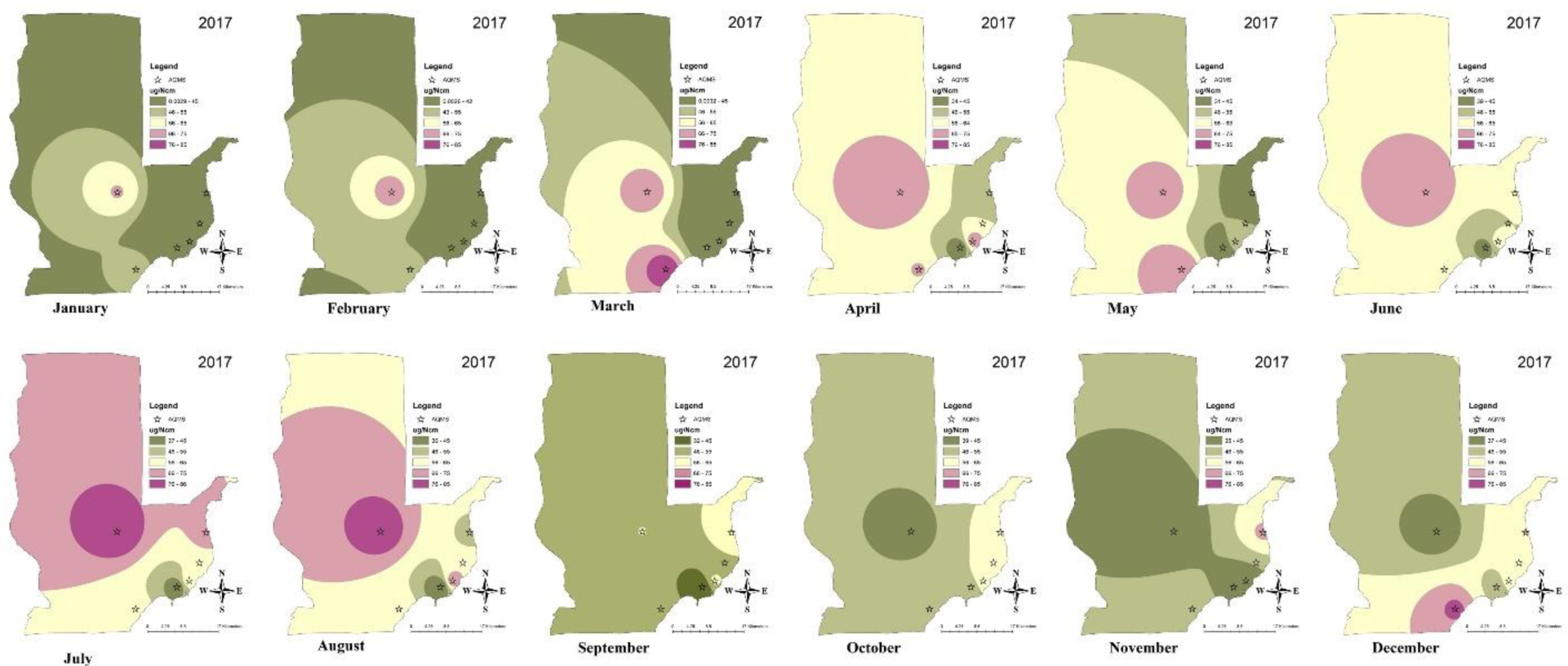
Particulate matter (PM_10_) concentration in the Davao City airshed in 2017 using the mean concentration (ug/Ncm) of each respective month from January to December.

PM_10_ distribution vary year to year. Most areas in all months during 2017 have a relatively PM_10_ concentration lower than 65 ug/Ncm concentration (Figure 5). At the same time, results show that the Davao City airshed experienced a high concentration of PM_10_ at 65-75 ug/Ncm throughout the year except September and October.

Areas close to DC Station 16 were exposed to this level from January to August which intensify to a critical concentration (75-85 ug/Ncm) and increased distribution during July and August. DC Station 14 have few months of PM_10_ levels at 75-85 ug/Ncm from March to May and in December.

PM_10_ distribution in 2018 do not resemble and was lower (< 45 ug/Ncm) than PM_10_ in 2017 (Figure 6). Fewer regions were at a critical level such as PM_10_ in DC Station 14 during May, July-August and DC Station 2 on June. PM_10_ concentration further decreased and improved the succeeding years. In 2019, all areas in the Davao City airshed were exposed to 65 ug/Ncm or lower and monthly distribution of PM_10_ varies (Figure 7).

**Figure 6.**
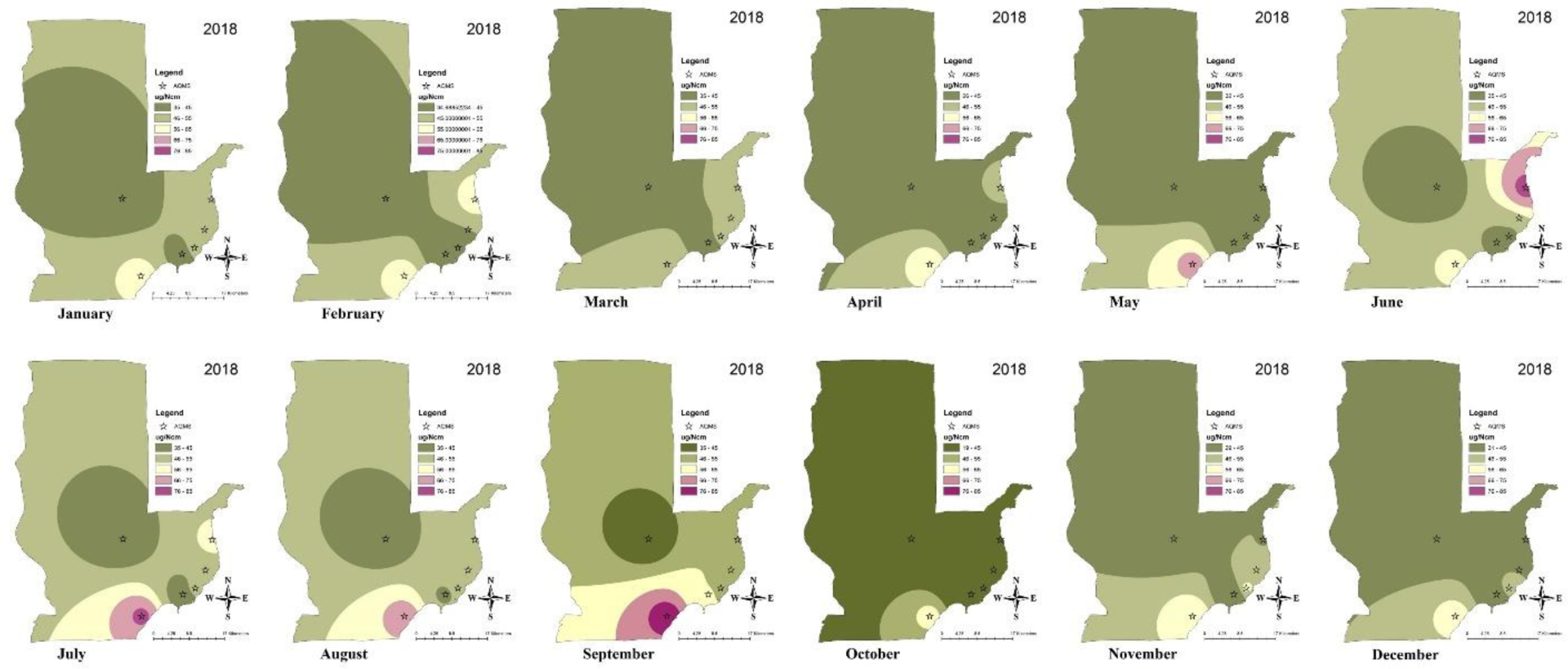
Particulate matter (PM_10_) concentration in the Davao City airshed in 2018 using the mean concentration (ug/Ncm) of each respective month from January to December.

**Figure 7.**
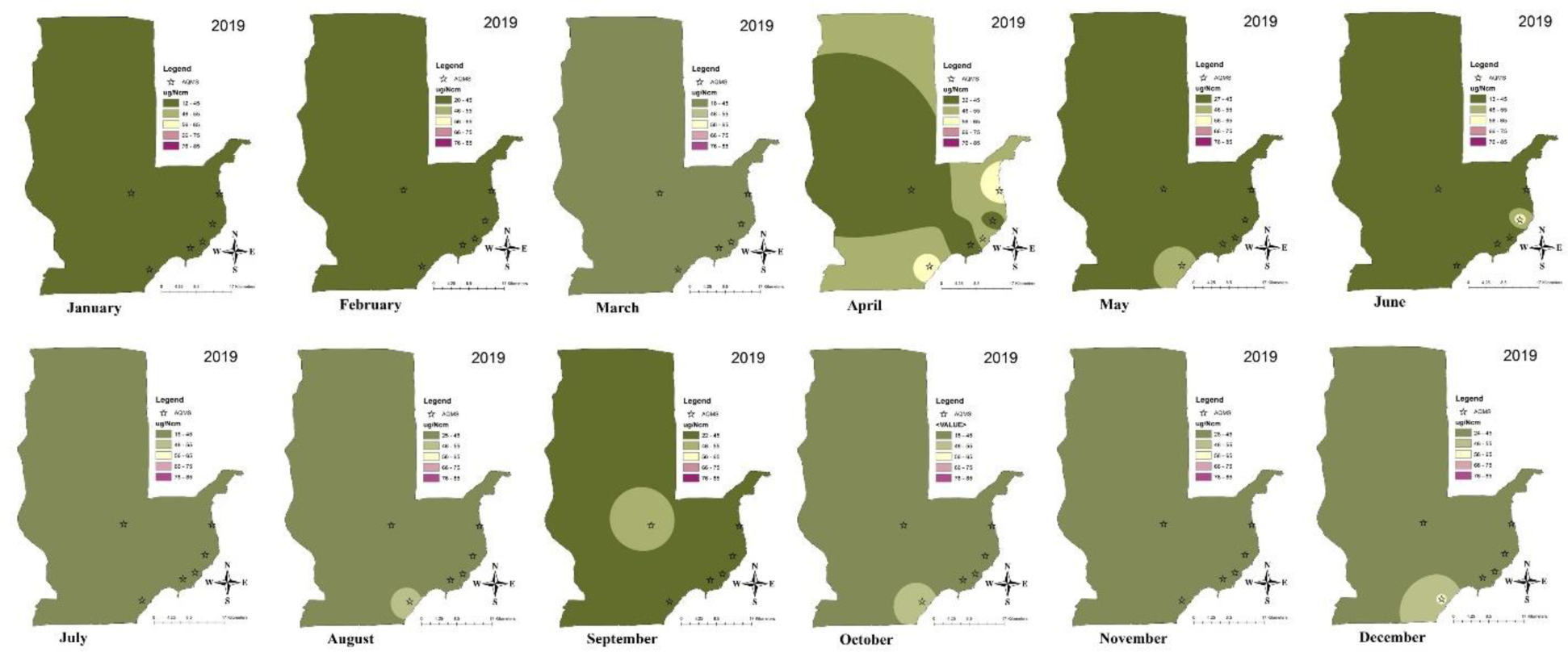
Particulate matter (PM_10_) concentration in the Davao City airshed in 2019 using the mean concentration (ug/Ncm) of each respective month from January to December.

Both PM_10_ distribution on 2020 (Figure 8) and 2021 (Figure 9) do not differ and PM_10_ concentration was lower than previous years with no seasonal variation observed. PM_10_ values was lower than 45 ug/Ncm although an evident high PM_10_ concentration was observed on April 2021. Apart from this difference, all months were exposed to PM_10_ concentration within the NAAQGV limit.

**Figure 8.**
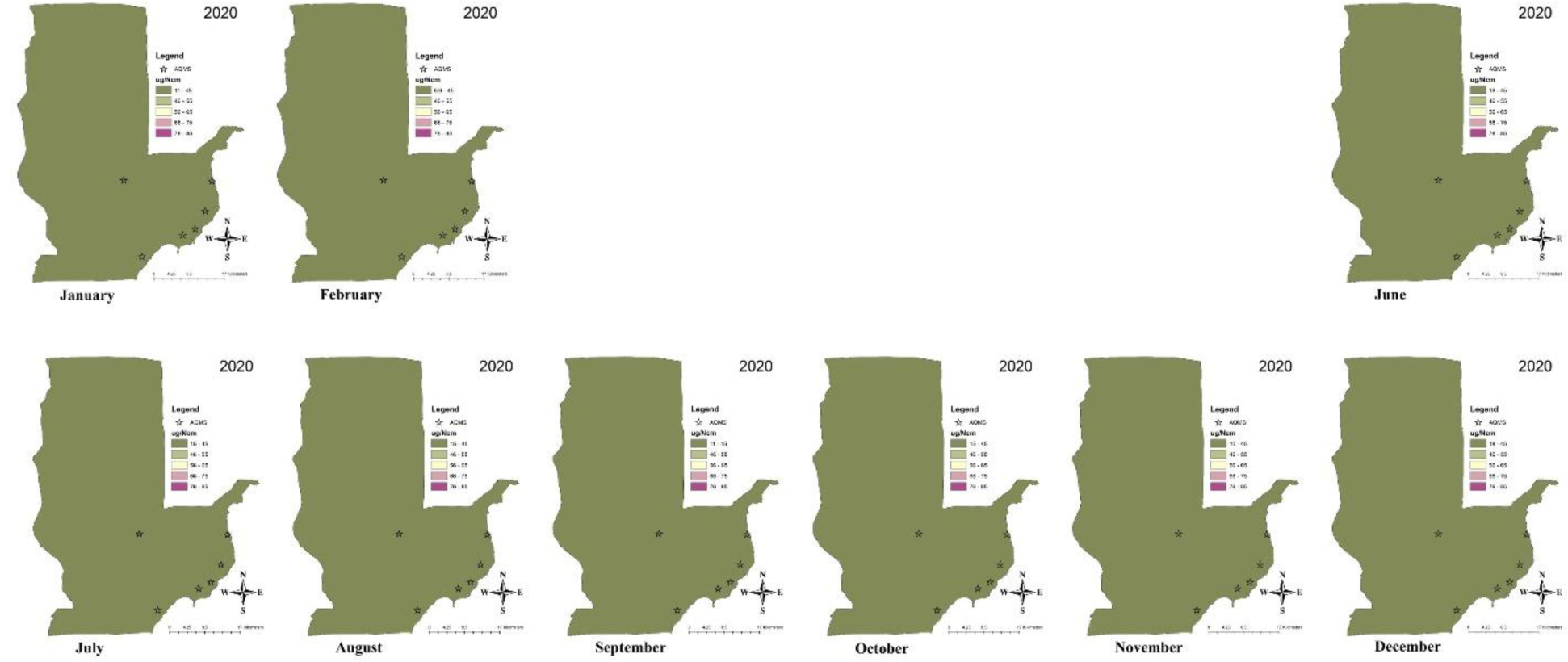
Particulate matter (PM_10_) concentration in the Davao City airshed in 2020 using the mean concentration (ug/Ncm) of each respective month from January to December.

**Figure 9.**
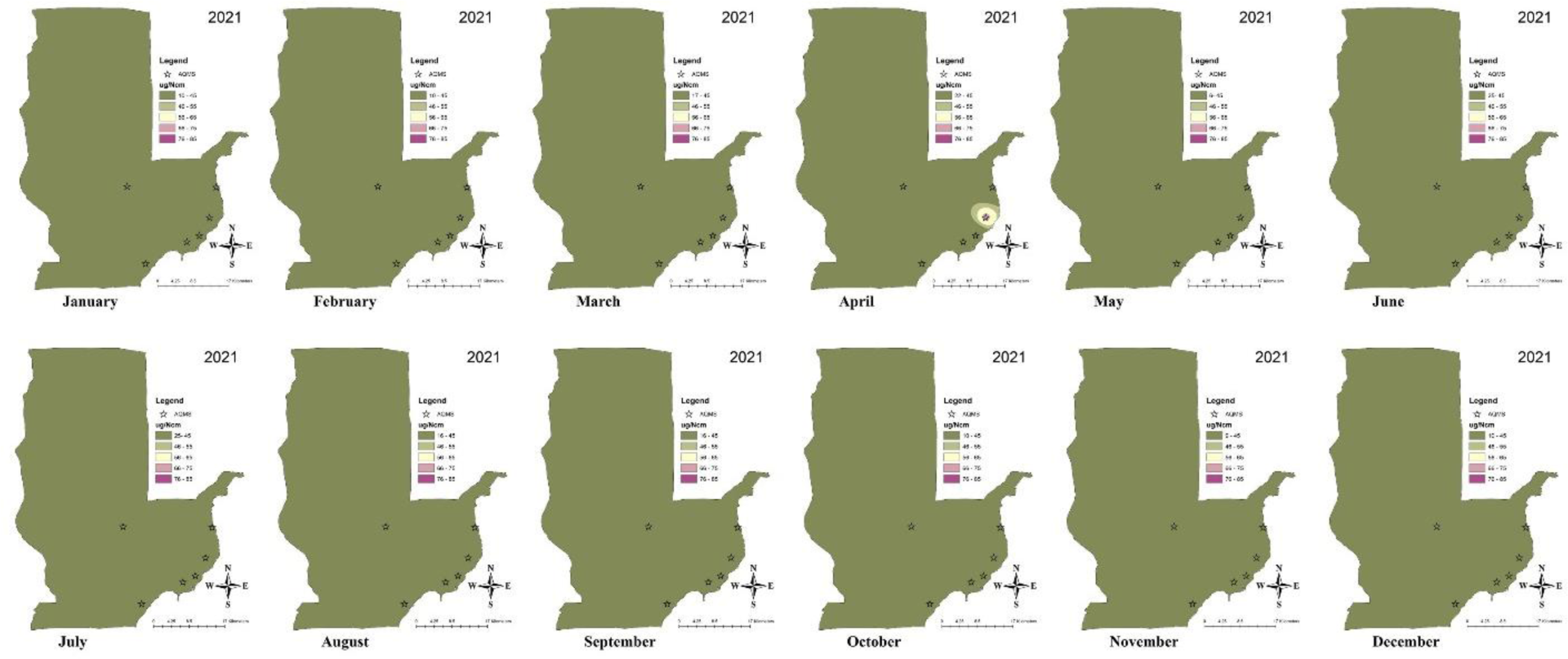
Particulate matter (PM_10_) concentration in the Davao City airshed in 2021 using the mean concentration (ug/Ncm) of each respective month from January to December.

Results show that reduction of anthropogenic emission sources (vehicle and industrial operation) in Northern India led to a decrease of air pollutants concentration such as carbon monoxide and different oxide compounds of nitrogen ^23^.

## 4. Conclusions

The six AQMS in the Davao City airshed provided air quality data essential to characterization of air quality status in the Davao City over span of years. Maps of PM concentration and distribution in Davao City reflected years of good and improving air quality conditions in the Davao City airshed. PM concentrations were categorized as good and within the limits of NAAQGV of the Philippines in 2016. The air quality data of the succeeding years from 2018 and onwards generally maintained “good” PM concentration level on both PMs (PM_10_ and PM_2.5_). An exception would be one AQMS location where successively for 3 years PM_2.5_ concentration was on an increased level (21 – 27 ug/Ncm). This variation was restricted to areas closer to one AQMS in the Davao City and it will be helpful to review activities not limited to emission sources but also social and recreational activities that may have contributed to this high PM_2.5_ level.

The availability of AQ data, in some instances, may be limited due to unexpected misfunction of AQMS samplers in some stations and may not represent the complete real-time concentration of air pollutants emitted in the region. Therefore, it is deemed important to secure or install more AQMS in the future for a more reliable and real-time measures of air quality in the city. This study using GIS technologies provided geostatistical and spatial data of air quality in the Davao City which are helpful tools (e.g. maps) not only in understanding previous and current air quality status in the region but also for environmental analysis of research scientists and decision making of air quality management regulatory agencies.

## Acknowledgements

The author sincerely thanks the Department of Environment and Natural Resources – Environmental Management Bureau XI, headed by the Regional Director Mario N. Bulacan and Engr. Liza Mae C. Villora for allowing the conduct of the study and the provision of air quality data. The author expresses her gratitude to DOST-PCIEERD for financial assistance and to Julie Atienza-Almasan for logistics support.

## Work Contribution

The author solely worked on the conceptualization, methodology development, software operation, and preparation of the manuscript.

## Funding

Financial support was provided by DOST-PCIEERD as part of the Balik Scientist Program.

## Notes

### Competing Interest Statement

The authors have declared no competing interest.

